# Determinants of the Transition Zone Width of Morphogen Readouts

**DOI:** 10.64898/2026.02.06.704367

**Authors:** Jan A. Adelmann, Roman Vetter, José M. Dias, Johan Ericson, Dagmar Iber

## Abstract

In tissue patterning, cell fate boundaries often form transition zones with mixed or gradual fates rather than sharp demarcations. Traditionally, such zones in morphogen-driven systems were attributed to fluctuations in morphogen concentrations, especially at low molecule numbers. Here, we present experimental data from the mouse neural tube, the precursor of the central nervous system, which challenges this view. Contrary to expectations, we find that the transition zone width (TZW) does not increase with distance from the morphogen source. By combining experiments, theory, and computational modelling, we identify key factors shaping the TZW. Our findings suggest that cellular readout noise, rather than morphogen fluctuations, determine the TZW. The inferred variability in the parameters defining morphogen dynamics and cellular readout remains within previously reported physiological ranges. This discovery adds to earlier findings that morphogen gradients maintain higher-than-expected patterning precision over long distances, offering new insights into the robustness of developmental processes.

## Introduction

As animals develop from a single cell, cells need to adopt the correct fate at the right position and time to form a functional organism. Morphogen gradients play a central role in establishing spatial boundaries. Virtually all simple morphogen gradients that have been quantified to date assume an exponential shape (Fig. 1A),

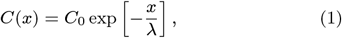

with an amplitude *C*_0_ at the morphogen source at *x* = 0 and a decay length *λ* [1–5]. According to the French flag model [6], concentration thresholds, *C*_*θ*_ = *C*(*x*_*θ*_), define boundary positions, *x*_*θ*_ (Fig. 1B). Cells exposed to morphogen concentrations above the threshold take on a different fate from those exposed to lower concentrations.

**Figure 1:**
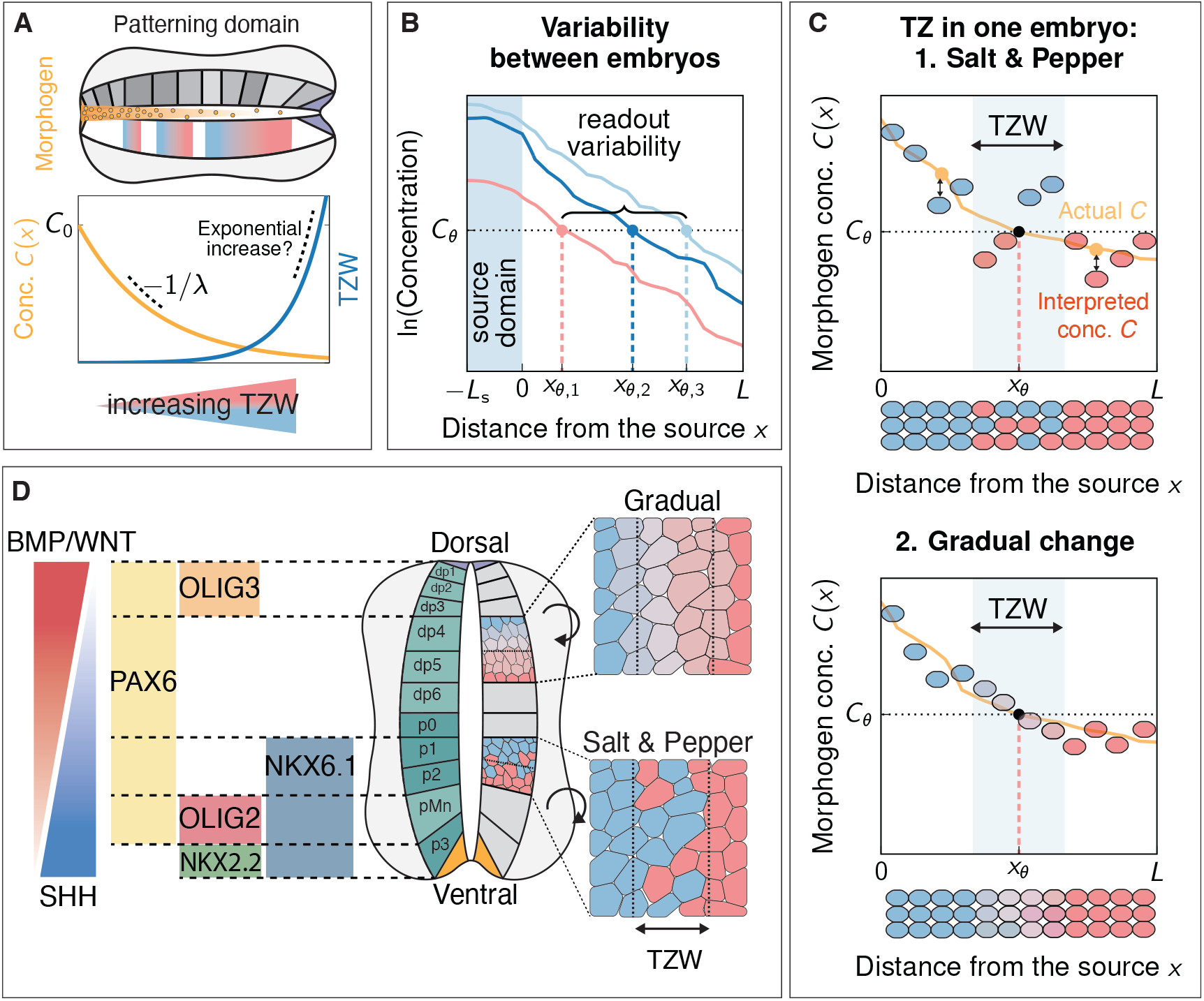
The positional error and the transition zone width in gradient-based patterning of the mouse NT. **A** SHH forms an exponential concentration gradient along the DV axis of the NT with amplitude *C*_0_ and decay length λ (orange line). The transition zone width (blue line) has been suggested to increase inversely proportional to the exponentially decreasing morphogen concentration due to low copy numbers. **B** Molecular noise in morphogen production, turn-over, and transport leads to different morphogen gradients in different embryos. As a consequence, the threshold concentration *C*_*θ*_ is reached at different positions *x*_*θ,j*_, leading to a positional error *σ*_*x*_. **C** At the boundary between readout domains, “salt and pepper” patterns (mixed cell fates, top) or gradual responses (bottom) are observed. **D** Patterning of the mouse neural tube by opposing SHH and BMP/WNT morphogen gradients. The gradients specify the location of progenitor domains characterised by a combination of distinct transcription factors. Not all domain boundaries are sharp. Instead, there are gradual transitions and “salt and pepper” patterns.

However, both morphogen gradients and the cellular readout processes are inherently noisy. As a result, the threshold concentration *C*_*θ*_ may be reached at different positions *x*_*θ,j*_ in different embryos, *j* (Fig. 1B). This inter-embryonic variability, the so-called positional error, can be quantified with the standard deviation of these readout positions [7],

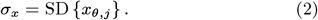

Beyond differences in readout position between embryos, a second source of imprecision concerns the sharpness of patterning domain boundaries (Fig. 1C). There is typically a transition zone at the boundary, where gene expression or cell fates are either mixed or change gradually. Traditionally, such transition zones have been attributed to local variability in morphogen concentration and “binding noise” in morphogen-receptor interactions, particularly at low molecule numbers. If morphogen levels fluctuate due to independent random events occurring with a probability proportional to time or space, their concentration can be described by Poisson statistics. For an exponential morphogen gradient (Eq. 1), the transition zone width (TZW) would then be expected to increase exponentially with its distance from the morphogen source [8] (Fig. 1A, see Supplementary Notes for details),

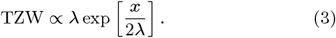

Such behaviour would severely limit the usefulness of morphogen gradients, as readout domains would become increasingly imprecise and fuzzy at larger distances from the source [8].

We recently developed a theoretical framework to estimate morphogen gradient precision based on molecular noise in morphogen production, decay, and transport rates [7]. In our simulations, we solved the steady–state diffusion equation for morphogen concentration on a cellular domain, where the production and degradation rates as well as the diffusion coefficient varied from cell to cell according to measured molecular noise levels. This analysis revealed that physiological noise results in far smaller differences in morphogen gradients between embryos than previously expected [7, 9–11]. Remarkably, the predicted positional error closely matches the reported small morphogen readout errors [7] and can reach subcellular precision across the physiological patterning field [11], suggesting that morphogen gradients should be sufficiently precise to support observed embryonic patterning accuracy [7, 12]. Further simulations revealed that small cell diameters and multiple parallel cell rows enhance patterning precision [9, 11], whereas previously suggested mechanisms such as self-enhanced morphogen degradation [13] play only a minor role [10]. In these simulations, the gradient profiles remained monotonic due to diffusion, precluding the emergence of a transition zone with mixed cell fates from fluctuations in the morphogen concentration, despite cell-to-cell differences in morphogen kinetics. This raises the key question: what determines the width of the transition zone?

The mouse neural tube (NT) has long served as a model system for studying gradient-dependent patterning and its precision [5, 12, 14–18]. The NT emerges along the back of the embryonic rostral-caudal body axis and gives rise to the central nervous system [19]. During NT development, distinct progenitor domains emerge along the dorsal-ventral (DV) axis, ultimately inducing different neuronal subtypes (Fig. 1D). The boundaries of the progenitor domains are defined by opposing exponential Sonic hedgehog (SHH) and bone morphogenetic protein (BMP)/WNT gradients (Fig. 1D). We now show that the TZW in the mouse NT is largely independent of the distance from the source, suggesting that it is not the Poissonian low molecule number limit that determines its size. We quantify the positional errors of several NT domain boundaries and find that they fall within previously reported ranges. Using our computational framework, we then infer molecular noise levels in morphogen production, decay, transport, and readout, again within physiological limits. In this way, we show that the TZW in the NT can be theoretically explained by cellular readout noise, while fluctuations in the morphogen gradient itself play no significant role. Both the positional error and the TZW remain small even far from the source, suggesting that morphogen gradients alone can achieve the observed patterning precision.

## Results

### Analysis of readout boundaries in the mouse NT

We analysed the expression patterns of five transcription factors (TFs), NKX2.2, OLIG2, PAX6, NKX6.1, and OLIG3, whose readout boundaries are spread along the entire DV axis of the mouse NT (Fig. 2A,B). We define the patterning domain to be the region between the dorsal end of the floor plate (FP) and the ventral end of the roof plate (RP). The readout position *x* of a domain boundary was measured either relative to the dorsal (D) end of the FP or the ventral (V) end of the RP (Fig. 2A). For each NT slice, the FP length, *L*_F_, was measured using a morphology-based approach, while the RP length, *L*_R_, was assumed to be equal to that of the FP (Fig. 2A).

**Figure 2:**
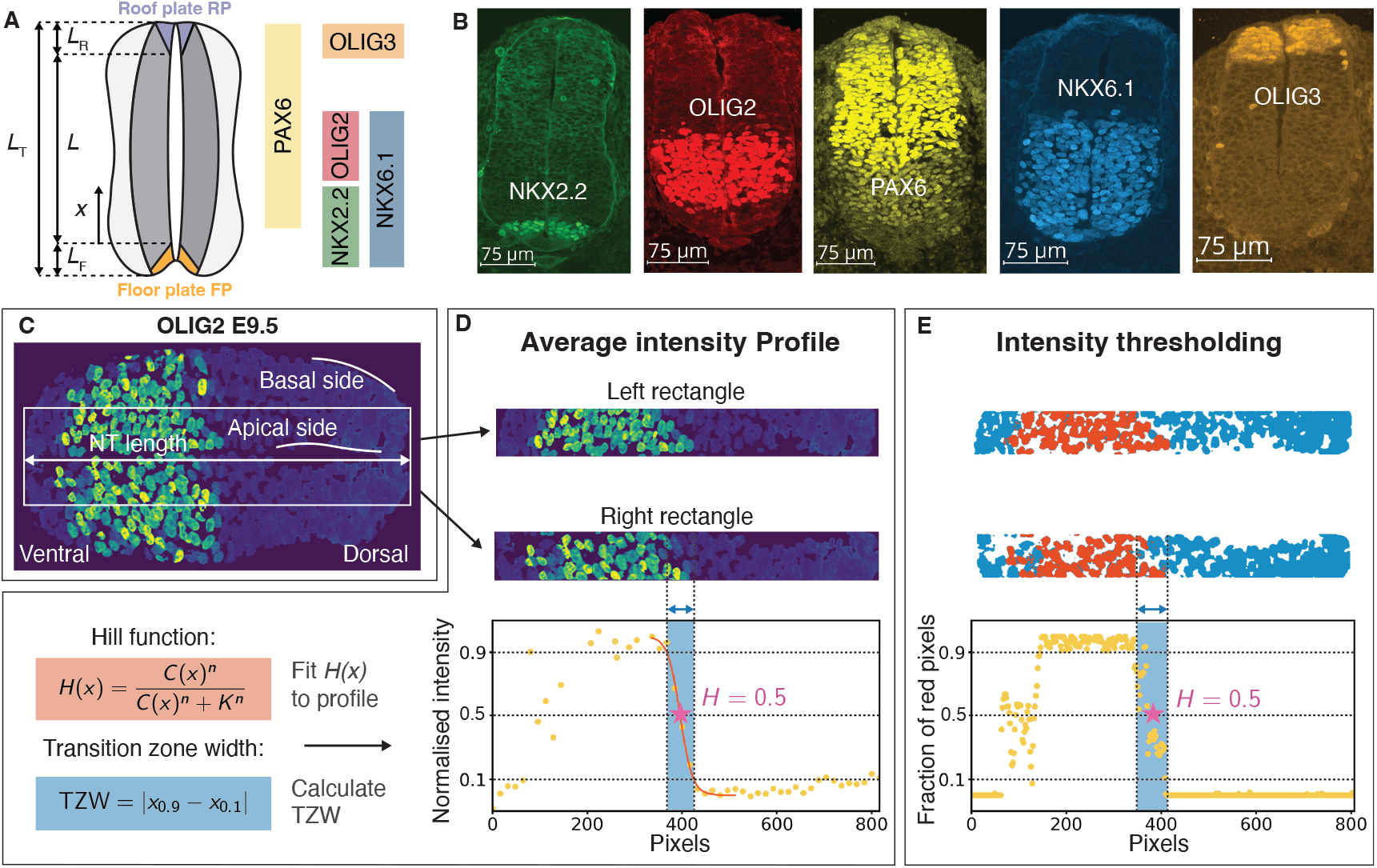
Workflow of progenitor domain analysis in the mouse NT. **A** Illustration of the coordinate system and patterning domains along the DV axis of the NT. **B** NT sections showing immunohistochemistry staining of the different boundaries analysed in this study. **C** Example NT section of OLIG2 at developmental stage E9.5. Each section is aligned with the *x*-axis and two rectangles are extracted along the apical tissue side. **D** Extraction of normalised intensity profiles form the rectangular regions. The intensity profiles were fitted with Hill functions, which enables the extraction of a transition zone width for each rectangle. **E** Intensity profiles extracted using multi-Otsu thresholding.

We used two different methods to determine the readout position *x* and to evaluate the positional errors and the TZWs (Fig. 2C–E; Methods). In the first, we fitted a Hill equation (Eq. 5) to the min-max normalised fluorescence intensity profiles and defined the TZW as the range over which the Hill curve decreases from 0.9 (90% normalised intensity) to 0.1 (10% normalised intensity) (Fig. 2D). The choice of this range is motivated by the fact that a broad interval offers an upper bound on the set of cells that continue to express the marker, thus most likely encompassing all cells within the corresponding domain. As an alternative method, we applied multi-Otsu thresholding to extract regions expressing the marker (Fig. 2D). We extracted a profile by counting the fraction of marker-expressing pixels (red) relative to background pixels (blue). This allowed us to define the TZW analogously as the range over which the expression transitions from 90% to 10%.

In both methods, we defined the readout position as the point where the profile crosses 0.5 (50%) (Fig. 2D,E), corresponding to the position *x*_*θ*_ where the morphogen gradient reaches the threshold concentration *C*_*θ*_.

We analysed data from three developmental time points, embryonic day (E) 9.5, 10.5, and 11.5 (somite stages 20–48), at the forelimb level (see Tables S1 and S2 for sample numbers). The NT grows substantially over time and the growth curve for somite stages 20–48 is linear (*R*^2^ = 0.89) [15]. This allows us to use the total NT length (*L*_T_) as a proxy for developmental time (Fig. 3A) [20]. Analysis of the data yielded similar readout positions for both approaches. For those progenitor domain boundaries that have been quantified before (NKX2.2, OLIG2, NKX6.1) [12, 14, 15, 18], we observe consistent positions (Fig. 3B). Any differences can likely be accounted to the different choice of intensity thresholds (50% vs. 10%).

**Figure 3:**
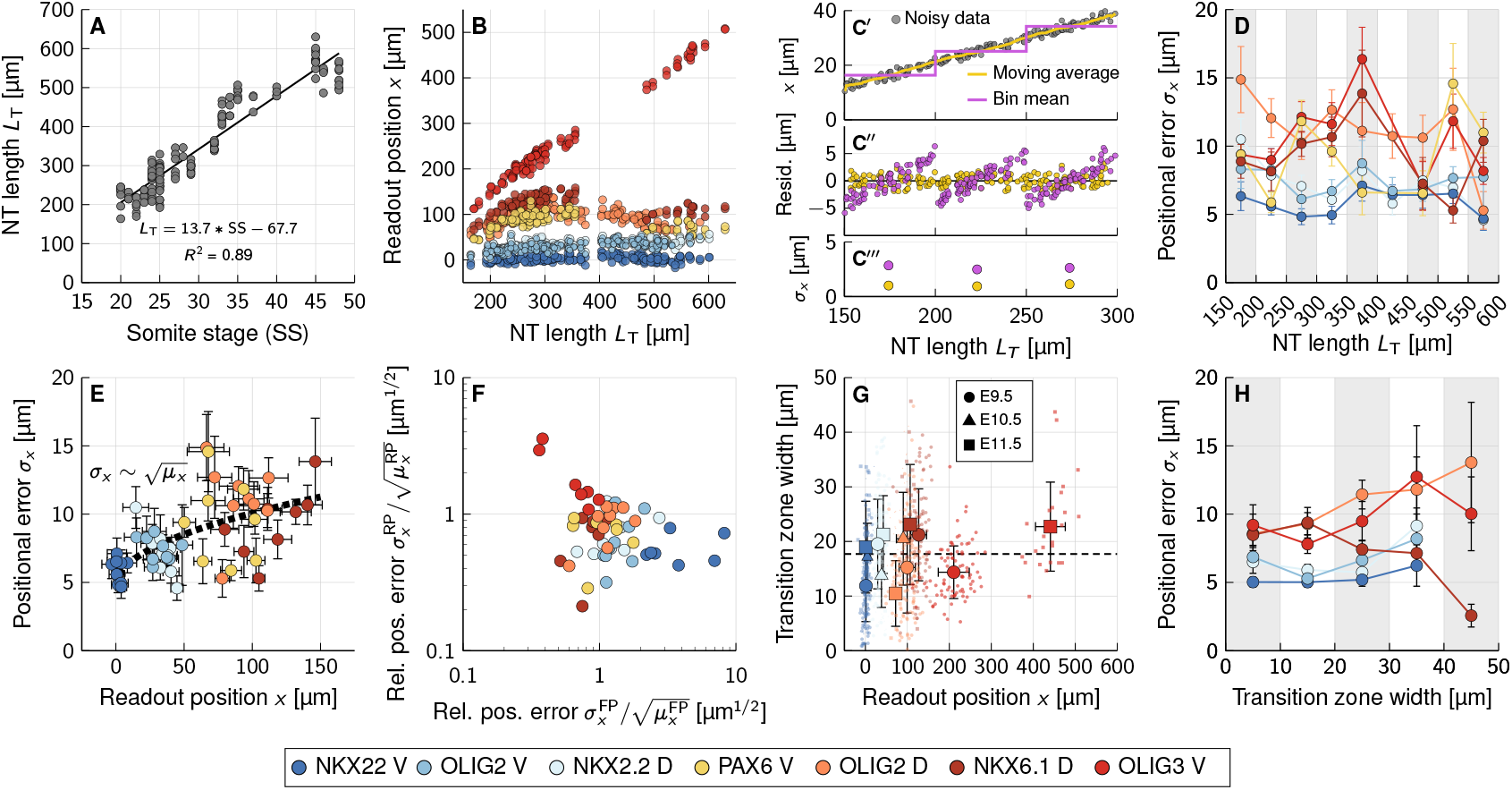
Quantification of positional errors and transition zones. **A** NT length over developmental time. **B** Progenitor domain boundary positions from the FP over developmental time, as measured by increasing total NT length *L*_T_. **C** Impact of temporal shifts on positional error estimation based on simulated data. **C**′ Simulated linear shift (grey dots) analysed by an adaptive moving average (yellow); 50 µm window, symmetrically truncated at boundaries) or a per-bin mean (magenta), which is stepwise and ignores within-bin motion. **C**″Residuals are symmetric and near zero for the moving average, while the per-bin residuals show systematic deviations over time. **C**‴ Per-bin mean overestimates *σ*_*x*_ (standard deviation of residuals) under temporal shifts. **D** Positional error over developmental time, calculated using a moving average windows of size 50 µm in *L*_T_ to avoid overestimation. **E** Positional error as in (D), vs. the boundary distance from the ventral limit of the FP. The black dotted line indicates the fitted theoretical scaling 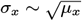 [9]. For fit parameters see Table S7. **F** Positional error measured relative to the dorsal end of the FP and ventral end of the RP, normalised by the square root of the mean readout position from the respective boundary. **G** The transition zone width (TZW) of the domain boundaries shows no systematic dependency on the readout position. Dashed line represents average TZW over all samples. **H** Transition zone widths vs. positional errors. TZWs were grouped into bins of length 10 µm. (B, D–H) All data points were obtained using Hill fits, except for the ventral OLIG3 boundary (thresholding method). For sample numbers in each bin, see Tables S1, S3, S4.

There are small but significant differences between the analysis approaches regarding the TZW. Generally, Otsu thresholding provides more accurate results for strong nuclear staining (e.g., OLIG3). However, it becomes less reliable for fading signals, as seen with OLIG2 at E11.5, when many progenitors lose the signal during differentiation. The dorsal OLIG2 boundary varies in shape, ranging from straight lines to V-shapes, which are better smoothed out by the Hill function fit than by thresholding, explaining the generally smaller TZW observed with the Hill fit. For the ventral boundary of OLIG3, the Hill fit overestimates the TZW due to challenges in capturing very narrow and sharp boundaries. Additionally, the gradual decrease in PAX6 intensity along the DV axis is poorly captured by thresholding, but better by the Hill fit. Due to variability in immunofluorescence staining and the strengths and weaknesses of each method, different boundaries are better quantified by one approach or the other. Moving forward, we will adopt the Hill function fit as the standard method, while using Otsu thresholding for the OLIG3 boundary. See Figs. S1–S3 for a comparison of the results.

### Quantifying the positional error in the mouse NT

Readout positions vary between embryos, resulting in a positional error *σ*_*x*_ (Eq. 2). As illustrated with synthetic data (Fig. 3C), tissue growth (Fig. 3A) and overall boundary movement (Fig. 3B) must be accounted for to avoid artificially inflating this error [7, 21, 22]. Calculating *σ*_*x*_ by simply binning data into 50 µm windows ignores within-bin trends, resulting in systematically biased residuals (Fig. 3C″) and a subsequent overestimation of the positional error (Fig. 3C‴). In contrast, an adaptive moving average centers the residuals around the local trend, isolating the underlying stochastic noise. Accordingly, we employed moving averages to eliminate these confounding temporal effects before grouping data into 50 µm bins for final error quantification (Fig. 3C′–C‴).

The positional error along the entire mouse NT remains largely below 15 µm (Fig. 3D) – roughly equivalent to three cell diameters [12, 15]. This aligns with previous measurements of NKX6.1, PAX7, and OLIG2 boundaries [12, 18]. No systematic change in positional error is observed over time (Fig. 3D), even though the distance of the boundary positions from the source changes (Fig. 3B). The positional error shows substantial scatter across bins, with larger uncertainty (error bars) in sparsely sampled bins (Fig. 3D, E). Nevertheless, the overall increase in positional error with readout position appears to be consistent with the theoretically predicted square-root scaling (black dotted line) [9].

Since embryos are grouped into bins within which the total NT length varies by at most 50 µm, the readout position is virtually equally variable between embryos whether it is measured from the ventral (FP) or dorsal (RP) limit (Fig. S3). Consequently, the positional error is almost identical when measured from either pole, ranging from approximately 5 to 25 µm. Within this symmetry, *σ*_*x*_ still increases with the square root of the distance from the respective reference boundary (Fig. 3E). However, once normalized as 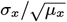, the values are very similar regardless of the side from which they are measured (Fig. 3F).

This robustness is particularly evident for OLIG3, a dorsal marker whose ventral boundary shifts away from the SHH source and approaches the BMP/WNT source as the DV axis extends (Fig. 3B). The ventral OLIG3 boundary moves from an average distance of 120 µm from the dorsal limit of the SHH source in the FP at E9.5 to about 500 µm at E11.5. Despite this increased separation from the FP, the positional error remains low and in a comparable range (5–15 µm) to estimates obtained when the same boundary is measured from the closer RP boundary (5–10 µm) with its BMP/WNT source (Fig. S3).

### The transition zone width is independent of the readout position

While the TZW of each analysed domain boundary varies widely between 0 and 50 µm (0–10 cell diameters) (Figs. 3G, S1E, S2), the average TZW across different markers and boundaries remains around 20 µm (4 cell diameters), independent of the readout position (Fig. 3G, dashed line) or developmental time point (for sample numbers see Table S1). This finding challenges the prevailing hypothesis that the TZW in the NT results from low morphogen concentration and should therefore increase exponentially with distance from the source [8].

The TZW and the positional error quantify different aspects of patterning precision. The positional error reflects inter-embryonic variability (location of domain boundary), whereas the TZW measures the spatial extent over which cells within a single embryo undergo a gradual or mixed fate transition. Consistent with this, we observe no strong correlation between TZW and positional error. The grouping of samples with similar TZW into 10 µm bins does not reveal higher positional errors in samples with larger TZW (Fig. 3H; see Table S4 for sample numbers in each bin), hinting at a partial decoupling of the mechanism controlling boundary placement accuracy and boundary sharpness.

The TZW of the ventral boundary of PAX6 is exceptionally wide (Fig. S2G,H). However, this is likely not the result of molecular noise, but of gene regulatory interactions. *Pax6* is repressed not only by NKX2.2, but also slightly less strongly by the more dorsally expressed OLIG2 [14]. In line with this, the ventral PAX6 boundary exhibits a gradual change in intensity compared to the sharp change of the other domains, and there is thus no salt and pepper pattern (Fig. 2B). The domain boundary is placed with the same accuracy as for the other domains, and the positional error of ventral PAX6 is thus low in comparison to its TZW (Fig. S2G,H).

These observations suggest that the TZW is not primarily set by fluctuations in the morphogen gradient itself, but rather by noise in the cellular readout process that converts morphogen concentration into gene expression. To test this idea, we modelled the readout of an exponential morphogen gradient with a Hill function, which provides a minimal description of a noisy, threshold-based decision process. In this framework, the TZW is defined as the spatial distance between two fixed relative readout levels, H_1_ and H_2_, reached at positions *x*_1_ and *x*_2_, respectively. For an exponential gradient, this yields

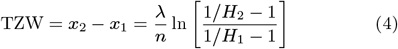

(see Supplementary Notes for a derivation). Strikingly, this TZW is independent of the readout position *x*_*θ*_ and thus of the local morphogen concentration. Instead, it depends linearly on the gradient decay length λ, and is inversely proportional to the sharpness of the readout, *n*.

In the mouse NT, we obtained a mean TZW close to the reported mean gradient decay length, *μ*_λ_ ≈ 20 µm [12] (Fig. 3G) when we determine the TZW as the range over which Hill functions decrease from *H*_1_ = 0.9 to *H*_2_ = 0.1 (Fig. 2D). Assuming that the thresholds of the noisy read-out process fall between *H*_1_ = 0.9 and *H*_2_ = 0.1, Eq. 4 requires an average Hill coefficient of *n* ≈ 4.4 to match the measured mean TZW. Excluding PAX6, our fits yield *n* = 3.4 ± 4.5 (mean ± SD), which is compatible with the predicted value, given the large uncertainty in inferring *n* from fits (see Table S8 for individual estimates). We conclude that the distance-independent TZW observed in the mouse neural tube can be explained as a geometric consequence of exponential gradients combined with a noisy but threshold-based cellular readout.

### The determinants of the transition zone width

Based on our analysis, the TZW appears to be independent of fluctuations in the morphogen concentration and rather seems to be determined by readout noise. To study the impact of readout noise on the TZW and positional error, we now extend our previous cell-based simulation framework of morphogen gradients (Fig. 4) [7, 9, 10].

**Figure 4:**
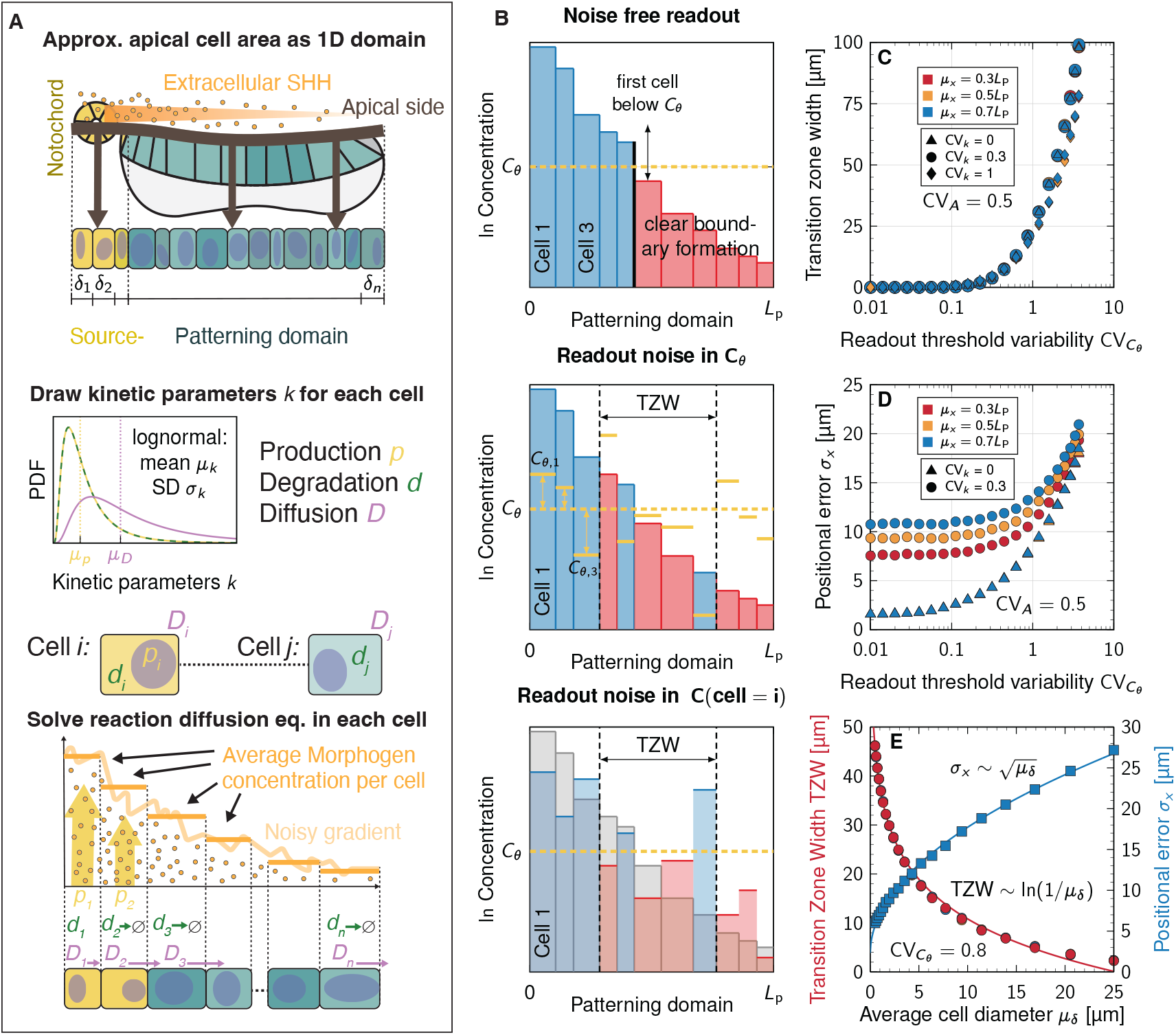
Simulation of noisy morphogen gradients. **A** The NT was modelled as as a one-dimensional cellular domain with source and patterning subdomains with average cell diameter *μ*_*δ*_ = 5 µm. Each cell has a set of unique kinetic parameters *k* = *p, d, D*, drawn from log-normal distributions with mean *μ*_*k*_ and standard deviation *σ*_*k*_. The steady-state reaction-diffusion problem was solved and the resulting morphogen concentrations were then averaged over each cell. **B** According to the French flag model, cells switch fate at threshold concentrations *C*_*θ*_. In a noise-free system, cells would switch fate at the correct position in the patterning domain (top panel). If cells sense an incorrect threshold concentration (yellow lines) they switch fate at the wrong position in the patterning domain and a transition zone of mixed cell fates emerges. This can either be due to biological variability in the threshold concentration *C*_*θ*_ (middle panel) or because cells read out a signal that deviates from the actual morphogen concentration (depicted as grey bars in the bottom panel). **C, D** Impact of readout noise levels on the TZW and positional error. **E** Impact of the cell size on the TZW (circular marks, red axis) and the positional error (square marks, blue axis). The shown logarithmic drop in the TZW is a phenomenological approximation. For fit parameters see Table S7. Data points are means from *N* = 10^4^ independent simulations, SEM are smaller than the marks.

SHH binds to its receptor PTCH1 on the cilium, which is restricted to the apical surface of the neuroepithelial cells (Fig. 4A, top row) [5]. Accordingly, we restrict our analysis to the apical surface along the DV axis, which we approximate by a 1D cellular domain. Note that the 1D approximation artificially increases the positional error compared to a 2D approximation [11]. The estimate that we obtain is therefore conservative and the expected positional error will be smaller. The readout of SHH is complex, given the restriction of the receptor PTCH1 to the cilium. However, we previously compared different morphogen readout modes and found that the restriction of the receptor to a cilium barely affects the readout precision in our model [9]. Accordingly, we average the concentration of the morphogen gradient over the entire 1D cell diameter, rather than explicitly representing the effect of the cilium (Fig. 4A, bottom row).

The variability in a model parameter between individual cells is expressed by its coefficient of variation, CV = *σ*/*μ*. For each cell, we draw an individual parameter value from probability distributions with a fixed mean and CV. In figures presenting the simulation results, unless stated otherwise, we report the sample CV to reflect the variability as it would be quantified from limited experimental data. For small and variable populations, the sample CV is biased and underestimates the true CV (Supplementary Notes).

Our model has the following parameters per cell: its diameter *δ*, the morphogen production rate *p*, decay rate *d*, diffusivity *D*, and the readout threshold *C*_*θ*_. As the morphogen production is restricted to the source, *p* is non-zero only in source cells, while *C*_*θ*_ applies only to cells in the patterning domain. Thus, all cells carry four stochastic parameters each. After setting up the source and patterning domains, a reaction-diffusion equation (Eq. 7) is solved with random cell-specific kinetic parameters *k* = *p, d, D*, drawn from log-normal distributions with fixed *μ*_*k*_ and CV_*k*_ (Fig. 4A, middle row).

The size of the morphogen-secreting source changes as NT development progresses, and it varies between embryos [15]. However, since the increase in source size over time, as well as its variability, do not a!ect the positional error developmental [9, 11] and the TZW for physiological noise levels (Fig. S4A–D), we set a fixed source size of 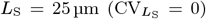 in the simulations.

The variability of the cell area, CV_*A*_, in the mouse NT has been shown to increase from 0.6 to 0.8 between E9.5 and E11.5 [23, 24]. As cell area variability in this range does not affect the positional error [9] and the TZW (Fig. S4E,F), we used the standard value of CV_*A*_ = 0.5 that we have used in our previous studies. The average apical cell area is fixed at a given stage, but decreases linearly from close to 10 µm^2^ at E8.5 to about 5 µm^2^ at E10.5 and E11.5 [24]. Cell areas are converted to cell diameters (Methods). The average cell diameter has previously been reported as *μ*_*δ*_ = 5 µm [5, 12].

Cell-to-cell variability in the kinetic parameters and the cell diameters results in differences in the morphogen profiles between embryos and thus a positional error that increases with CV_*k*_ (Fig. S4G) [7]. The small positional error for low kinetic noise levels is due to cell area variability but the impact at physiological levels is negligible (Fig. S4G). Despite variability in the kinetic parameters and cell diameters, the simulated morphogen gradients decrease monotonically, such that the fate transition is sharp (Fig. 4B, top row). To achieve the formation of a TZ in our simulations, we introduce noise in the readout of the morphogen gradient. Noise in the readout can, in principle, occur at two levels: Cells either use variable concentration thresholds *C*_*θ*_ (Fig. 4B, middle row), or they detect an imprecise morphogen concentration (Fig. 4B, bottom row). In our simulations, we do not distinguish between these types of error, and combine their effects into a single parameter, termed readout noise, a cell-specific threshold with variability 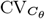.

Readout noise causes cells to switch fate at the “wrong” signalling level, leading to the emergence of a TZ that widens as 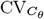 increases (Fig. 4C). In the simulations, the TZW is solely defined via concentration thresholds *C*_*θ*_. In agreement with the experimental data and the analytical solution for the Hill readout, we again find no dependency on the readout position (same symbols but different colours overlap in Fig. 4C). At moderate kinetic noise levels (CV_*k*_ = 0.3, circular marks), the TZW exhibits the same qualitative increase as in simulations without kinetic noise (triangular marks). Only at high levels of both kinetic and readout noise does kinetic noise impact the TZW. Interestingly, under these conditions, the TZ narrows, but still remains independent of readout positions. This small effect arises from the simulated gradients having sharp intermediate drops at large kinetic noise levels, resulting in shorter TZs (Fig. 4C, CV_*k*_ = 1, diamond marks).

In contrast, the positional error is predominantly determined by kinetic noise when readout noise is low (Fig. 4D). Unlike kinetic noise, readout noise does not exhibit spatial dependence (triangular marks from different readout positions overlap). The positional error remains largely constant at low readout noise levels, but increases substantially at high levels. At low readout noise levels, the positional error is dominated by kinetic noise (circular marks CV_*k*_ = 0.3 vs. triangular marks CV_*k*_ = 0).

In previous work, we demonstrated that the positional error increases with the square root of the cell diameter in the domain [9]. This scaling also holds with readout noise (Fig. 4E, blue, Fig. S4H). The TZW shows opposite behaviour: the larger the cells, the smaller the TZW, and there is no dependency on the readout position (Fig. 4E, red). The average TZW of 20 µm found in our data analysis (Fig. S2A–F) corresponds to an average cell diameter of approximately 4–5 µm (Fig. 4E). This aligns well with the cell diameter reported for the mouse NT [5, 12], which in turn corresponds to a positional error of approximately 10 µm or two cell diameters.

In summary, noise in morphogen production, degradation, transport, as well as in the cell size and the source size affects the inter-embryonic variability, i.e., the positional error, but not the sharpness of the domain boundaries, i.e., the TZW—at least not within the physiological range. Noise in the cellular readout, on the other hand, affects primarily the TZW and to a lesser extent the positional error. Finally, a larger average cell diameter increases the absolute positional error and reduces the TZW, possibly explaining why the apical cell diameter has not shrunk further in the evolution of patterned tissues [9].

### Inference of kinetic and readout noise levels

Building on the preceding data analysis, which estimated readout positions, positional errors, and TZWs across various progenitor domain boundaries in the developing mouse NT, we now apply our computational framework to infer the levels of kinetic and readout noise necessary to account for the observed positional errors and TZWs. Specifically, we used the average positional error *σ*_*x*_, the average TZW, and the average readout position *μ*_*x*_ of each domain as input to the gradient simulations.

We performed parameter screens across different levels of readout noise (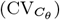) and molecular noise (CV_*k*_) (Fig. 5A,B). For each parameter combination, we obtained a corresponding positional error and TZW at a given readout position within the tissue, allowing us to plot contour lines that match the positional error and TZW obtained from the data (black lines in Fig. 5A–D). Their intersection point identifies the parameter combination that reproduces the observed values in the NT (Fig. 5B).

**Figure 5:**
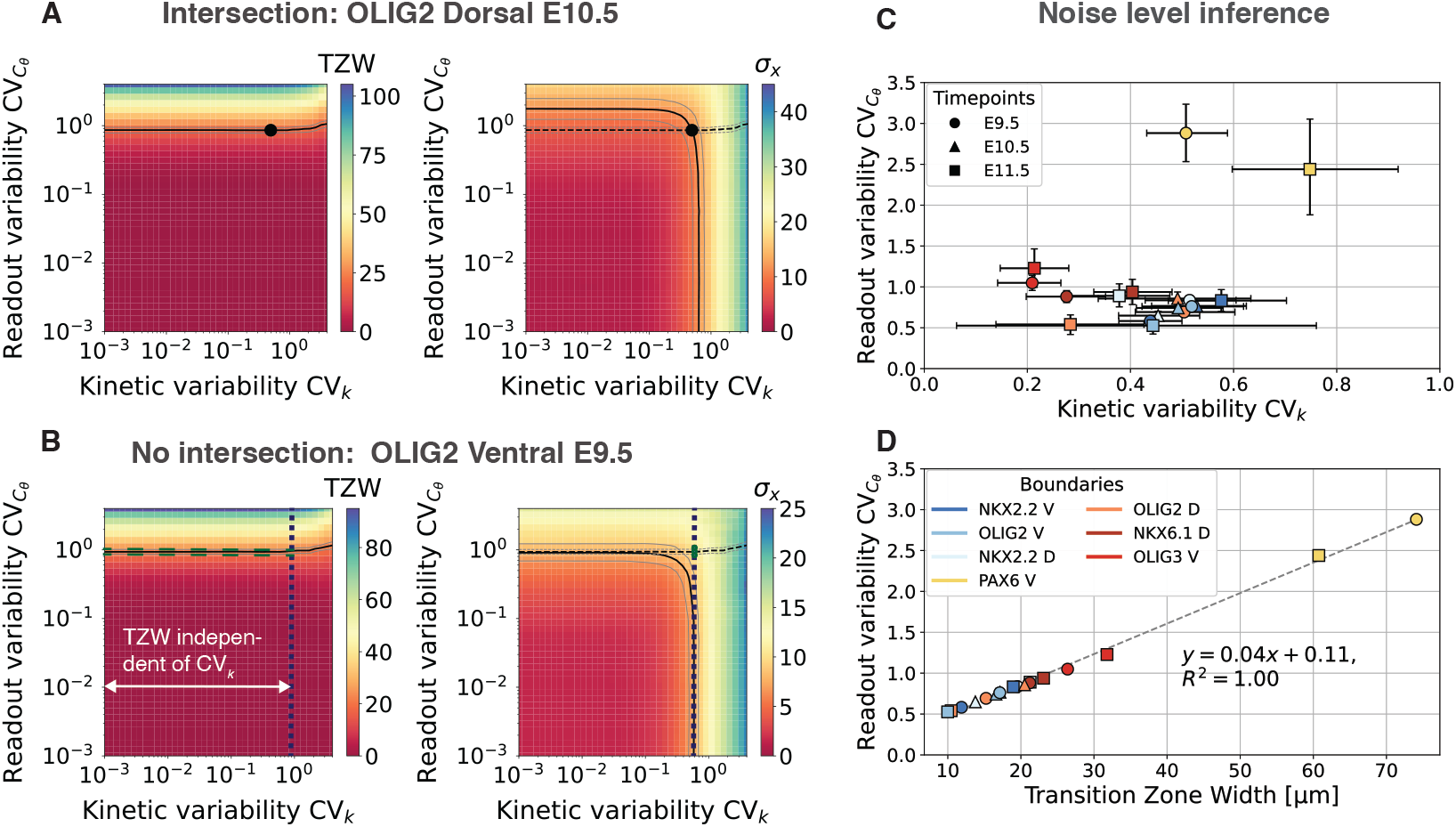
Inference of kinetic variability and readout noise. **A**,**B** Impact of noise levels in the morphogen kinetics and readout process on the TZW and the positional error, measured in µm. Each coloured square represents the mean over *N* = 10^4^ independent simulations. Contours correspond to mean TZW ± SEM or mean *σ*_*x*_ ± SE M. Exemplary results obtained from the Hill fit approach are shown; see Figs. S5 and S6 for individual plots. **C** Inferred noise levels (contour intersection points) for the different domain boundaries in the mouse neural tube. Error bars are SEM. **D** Linear relationship between the measured TZW and inferred readout noise level 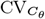. Error bars are omitted for clarity

For the contours to intersect, the TZW and positional error must be of similar magnitude (Fig. 5A). In some domains, however, the TZW is slightly too large for an intersection to occur (Fig. 5B). In these cases, we employ a conservative approach to estimate an upper bound for the noise levels. We select the maximum observed CV_*k*_ along the contour line, combined with the 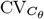 value at the intersection of the 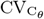 contour with the vertical line at max(CV_*k*_) (Fig. 5B, right panel, green line). In the tested range CV_k_ < 1, kinetic variability does not impact the TZW, making the intersection a good estimate for readout noise levels.

Note that uncertainty in the TZW, represented by the standard error in Fig. 5, may underestimate the actual readout noise variability. For consistency, we use standard errors to represent both TZW and positional error uncertainties, allowing for a direct comparison between the two measures.

Estimates of molecular noise levels are primarily available from measurements in the *Drosophila* wing disc. For Hedgehog (Hh), the ortholog of mouse SHH, the CV of the diffusion coefficient has been measured as CV_*D*_ = 0.18 [25]. Additionally, for Decapentaplegic (Dpp), the ortholog of mouse BMP4, the following values have been reported: CV_*d*_ = 0.5, CV_*p*_ = 0.59, and CV_*D*_ = 0.5 [26]. Our analysis shows that the kinetic noise levels of CV_*k*_ 0.2–0.6 fall within the previously reported ranges, with no apparent correlation with developmental time. The actual kinetic variability is likely lower than our estimates, as we adopt a conservative approach for boundaries with no intersection. In such cases, any value smaller than the reported CV_*k*_ is possible. Similarly, the readout variability 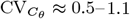 (excluding ventral PAX6) does not show correlation with developmental time either (Fig. 5C).

Our model does not account for regulatory interactions. Wide TZW that are a consequence of regulatory networks, rather than readout noise, are therefore not captured. This limitation is evident in the ventral PAX6 domain boundary, where kinetic noise levels appear significantly higher than otherwise (Fig. 5C). Boundary placement is as precise as in other domain boundaries, as the kinetic noise levels are within the range of the other domain boundaries. See Fig. S5 for the complete set of parameters screens with the Hill fit method and Fig. S6 with the thresholding method.

The measured TZWs and the readout noise levels inferred from the parameter screen are linearly related (Fig. 5D).

## Discussion

Here, we provide the first quantitative characterisation of transition zone widths for multiple progenitor domain boundaries along the dorsal-ventral axis of the mouse neural tube. We found that the width of the transition zone between progenitor domains is independent of their DV position. This absence of a positional dependency, particularly the observation that the transition zones do not widen with distance from the morphogen source, suggests that there is no substantial influence of “binding noise” in the patterning process of the NT that would result from low receptor occupancy, as previously hypothesised [8]. Instead, our results suggest that ligand numbers remain sufficient throughout the patterning process, avoiding a Poissonian limit, and allowing morphogen gradients to reliably instruct positional identities across the neural tube [7, 9–11, 20]. This is further supported by simulations that demonstrate that the TZW does not depend on the readout position but only on the readout noise levels. These simulation results are not limited to the NT but should also be applicable to others systems patterned by morphogen gradients.

We distinguish between the TZW and patterning precision, quantified by the positional error. While both describe aspects of boundary definition, they capture fundamentally different processes: The TZW reflects the spatial sharpness of the boundary, whereas the positional error measures the reproducibility of boundary placement across embryos. Our approach therefore provides a framework to quantify both aspects in parallel and to disentangle their contributions in the NT. By comparing the observed TZWs to our theoretical modelling framework, we have indirectly estimated the variability of the readout process, which might become accessible to experimental validation one day.

A particularly intriguing outcome of our analysis is the relationship between TZW and positional error as a function of cell size. Our simulations predict that larger cells lead to sharper boundaries (Fig. 4E), whereas smaller cells lower the positional error [9]. These opposing trends suggest that the in vivo cell size may represent a sweet spot that balances the need for sharp boundaries with the need for precise positioning, and might be the result of an evolutionary optimisation. Such a trade-off may be a general design principle of morphogen-patterned tissues.

In analysing the TZW, we assumed that the readout position is predominantly determined by the local morphogen concentration and the cellular readout process. As illustrated by the ventral PAX6 boundary, this is not always the case. The wide, gradual intensity transition at the ventral PAX6 boundary is likely the consequence of gene regulation rather than molecular noise. Specifically, NKX2.2 and OLIG2 both repress *Pax6* expression, with the more dorsal factor, OLIG2, having a weaker impact [14]. Regulatory feedbacks likely also impact other transition zones, though not as strongly as PAX6. In addition, cellular readout processes can introduce substantial levels of noise, or reduce noise by spatial and/or temporal averaging [1, 27–31]. Boundary sharpness can also be increased via mutual repression in transcription factor networks [14, 18], and cell sorting [32, 33]. Notwithstanding these many confounding factors, our finding of a distance-independent TZW rules out a dominant effect of binding noise on the TZW.

Finally, we note that estimating the TZW from DV slices may overestimate both the positional error and the TZW due the pseudo-stratification of the neural tube. Since the readout of SHH is limited to the apical surface, the pseudo-stratified organisation of the neural tube could artificially inflate the TZW and positional error measurements based solely on the DV axis [34]. 3D simulations involving irregularly shaped pseudostratified epithelial cells [34–36] may reveal more complex concentration profiles than those captured in the current simulation framework. Therefore, it remains to be explored whether the current approximations underestimate the precision of domain boundary formation. To address this, future studies may need to incorporate fluorescence sections from the apical surface of the neural tube in conjunction with 3D stochastic cell-based morphogen gradient simulations to fully capture the precision of morphogen-gradient-based patterning.

In summary, our results reveal that morphogen-based patterning in the NT is more precise and robust than previously anticipated. By quantifying both TZW and positional error, and linking them to cell size, we identify a potential evolutionary trade-off that may optimise tissue patterning.

## Methods

### Embryo collection

Mouse embryos were collected at different embryonic days (E9.5– E11.5) and staged by somite number. Mouse embryos were fixed in 4% paraformaldehyde (PFA) for 1.5 hours on ice with shaking. They were subsequently washed once with phosphatebuffered saline (PBS), cryoprotected by equilibration overnight in 30% sucrose in PBS, embedded in O.C.T., frozen on dry ice, and cryostat-sectioned in the transverse plane at 12 µm. Tissue sections were stored at − 80°C or directly analysed by immunohistochemistry.

### Immunohistochemistry

Immunohistochemistry of the tissue sections was carried out in a water-humidified chamber. The tissue on slides was incubated with blocking solution (3% FBS/0.1% Triton-X100 in PBS) at room temperature (RT) for 1 hour followed by incubation with primary antibodies overnight at 4°C and fluorophore-conjugated secondary antibodies for 1 hour at RT. Both primary and fluorophore-conjugated secondary antibodies were diluted in blocking solution. Sections were mounted using Vectashield (Vector Laboratories, Inc.) and coverslipped for imaging. Primary antibodies used: mouse anti-NKX2.2 (74.5A5), NKX6.1 (F65A2), PAX6 from Developmental Studies Hybridoma Bank (DSHB); rabbit anti-OLIG2 (AB9610, Millipore), and rabbit anti-OLIG3 (C. Birchmeier). For each position along the forelimb region, three images covering the entire length of the neural tube and separated by 1 µm were acquired using a Zeiss LSM880 confocal microscope, and a maximum intensity projection image was generated and used for subsequent analysis.

### Extraction of intensity profiles

The different progenitor domains in the mouse NT are marked by a combination of distinct transcription factors (TFs), in this work we focus on NKX2.2, OLIG2, NKX6.1, and PAX6 (Fig. 2A). From each image we extracted the neural tube using hand-drawn masks. The extracted neural tubes were rotated, using the apical side as reference, such that the DV axis aligns with the *x*-axis (Fig. 2B). To increase the number of samples, we positioned rectangles with widths of 100 pixels for developmental timepoints E9.5 and E10.5, and 50 pixels for timepoint E11.5, around the apical side of the neurothelium, creating ‘left’ and ‘right’ rectangles (Fig. 2B–D). Marker intensities within each rectangle were extracted using nuclear masks, created from the DAPI channel by Otus’s thresholding method. We extracted marker profiles using two different approaches. In both approaches we first divided the DV axis into 64 bins, each 16

pixels wide. In the first approach, we computed the average intensity in each bin and min-max normalised the result (Fig. 2C). Second, using multi-Otsu thresholding we segmented the intensity images to extract the regions expressing a specific maker (Fig. 2D). Next, we counted the number of positive pixels (pixels expressing the marker, red in Fig. 2D) and normalised by the total number of pixels in each bin to get an expression profile.

### Analysis of the transition zone width

Based on the marker profiles, we computed the transition zone width using two different methods:

#### 1. Hill fit method

Intensity profiles of the different domains were extracted and fitted to a Hill equation of the following form (Fig. 2C):

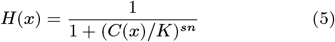

where the sign *s* in the exponent switches between a positive and a negative Hill function to account for the two boundaries per expression domain:

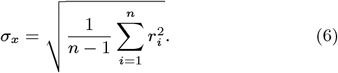

*C*(*x*) = exp[−*x*/λ], represents the exponentially decaying gradient profile with *x* = 0 at the ventral source, λ is the gradient decay length set to 20 µm, *K* the half-occupation constant, and *n* the Hill coefficient. By inverting Eq. 5 and solving for two different thresholds, here H(*x*_0.9_) = 0.9 and H(*x*_0.1_) = 0.1, the transition zone width is defined according to TZW = *x*_0.9_ − *x*_0.1_ (Fig. 2C). The readout position is defined to lie at the threshold H(*x*) = 0.5, where *K* = *C*_*θ*_.

#### Thresholding method

We applied multi-Otsu thresholding [37] to the NT sections. Pixels below the threshold were treated as not sufficiently expressing the marker. We then defined the transition zone as the region where the fraction of pixels labelled as expressing the TF in a pixel row perpendicular to the patterning axis drops from 90% to 10%. The readout position was set to be where this fraction crosses 50%. If a threshold value (90%, 50%, 10%) was crossed more than once, the median position was taken.

### Positional error calculations

The positional error *σ*_*x*_ at a given developmental stage is the standard deviation of the readout positions between embryos at that stage. To accurately estimate it, it is necessary to distinguish between variability in the readout position due to noise in the patterning process and variability caused by tissue expansion, which shifts the domain boundaries. The latter does not contribute to patterning inaccuracy as quantified here, and leads to overestimation of the positional error if not factored out [7]. To disentangle these aspects, we implemented a moving average filter with a window width of *w* = 50 µm, using the NT length along the DV axis as a proxy for developmental time (Fig. 3A, C′–C‴). If a sample *i* has NT length *l*_*i*_, we computed the moving average readout position 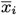 as the average readout positions of all samples whose NT length lies within the interval [*l*_*i*_ − *w*/2, *l*_*i*_ + *w*/2]. The residual 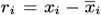 represents the deviation of a sample from the expected readout position at its stage. The positional error in a bin with *n* samples in it can then be computed as the standard deviation about the local moving average in that bin:

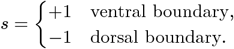

Repeating this process for all stage bins yields the positional error over developmental time. Note that to avoid biases in the first and last bin, the averaging window is shrunk to the distance to the first or last data point. The first and last residual, which then both vanish, are excluded from the calculation in Eq. 6. This method effectively removes any artificial positional error from the calculation that arises from different patterning domain lengths within a bin. The bin width can then be chosen freely without affecting the accuracy of the estimate much, as long as *n* remains sufficiently large. We set it equal to the moving average window here (50 µm), such that the number of samples in each stage bin, during which the NT grows by 50 µm, is about *n* = 20 on average in our dataset.

### Simulation of stochastic morphogen gradients

To obtain noisy steady-state morphogen gradients, we solved a stochastic reaction-diffusion equation for the morphogen concentration *C*(*x*) (Eq. 7) on a 1D cellular domain, *x* [*L*_S_, *L*_P_], as done previously [7]:

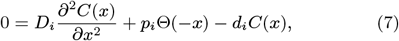

where *i* denotes the index of the *i*-th cell in the domain, and Θ is the Heaviside step function. The kinetic parameters *p* (morphogen production rate), *d* (degradation rate) and *D* (diffusion coefficient) were assumed to vary independently from cell to cell. Zero-flux boundary conditions were imposed at both tissue endpoints, assuming absence of a net morphogen flux in or out of the neural tube [20]:

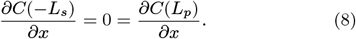

Cell areas *A*, production rates *p*, degradation rates *d* and diffusivities *D* were drawn independently from log-normal distributions. In a previous publication, we demonstrated that as long as the probability distributions meet certain criteria, the simulation results are largely independent of the specific distribution assumed [9].

The simulations proceeded as follows: First, the source and patterning domains of length *L*_S_ = 25 µm and *L*_P_ = 625 µm, respectively, were constructed by adding cells of diameter 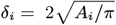 until the target lengths were matched. To obtain an average cell diameter of *μ*_*δ*_ = 5 µm with the desired log-normal cell area distribution with coefficient of variation CV_*A*_, we set 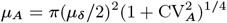 as derived in [9]. Mean parameter values were set to 20 µm.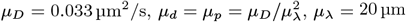

Next, the kinetic parameters *p*_*i*_, *d*_*i*_ and *D*_*i*_ were drawn for each cell, and Eq. 7 was solved using Matlab’s fourth-order boundary value problem solver bvp4c. Subsequently, the average concentration was calculated in each cell. All simulations were repeated *N* = 10^4^ times with independent parameters. Noise in the morphogen readout process was introduced by drawing the readout threshold *C*_*θ*_ from log-normal distributions for each cell.

Unless specified otherwise, parameters were fixed to the following values: *L*_S_ = 25 µm, *L*_P_ = 625 µm, CV_*A*_ = 0.5, 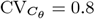, and CV_*k*_ = 0.3.

## Code Availability

The source code is released under the 3-clause BSD license. It is available as a public git repository at https://git.bsse.ethz.ch/iber/Publications/2026_adelmann_transition_zone.

## Acknowledgements

We thank Marco Meer and Liepa Kazlauskaitė for preliminary work and C. Birchmeier for rabbit anti-OLIG3. This work was funded by the Knut and Alice Wallenberg Foundation (KAW2011.0661, KAW2012.0101), the Swedish Research Council (2022-01217) and Hjärnfonden (F023-0035) to JE.

## Author Contributions

DI and JE conceived the study. JMD obtained the data. JA processed and analysed the imaging data, wrote the code, performed the simulations and produced the figures. JA, RV and DI analysed the results and wrote the manuscript.

