## Supplemental text for "Determinants of the Transition Zone Width of Morphogen Readouts"

### Determinants of the Transition Zone Width of Morphogen Readouts: Supplementary Information

#### 1 Supplementary Figures

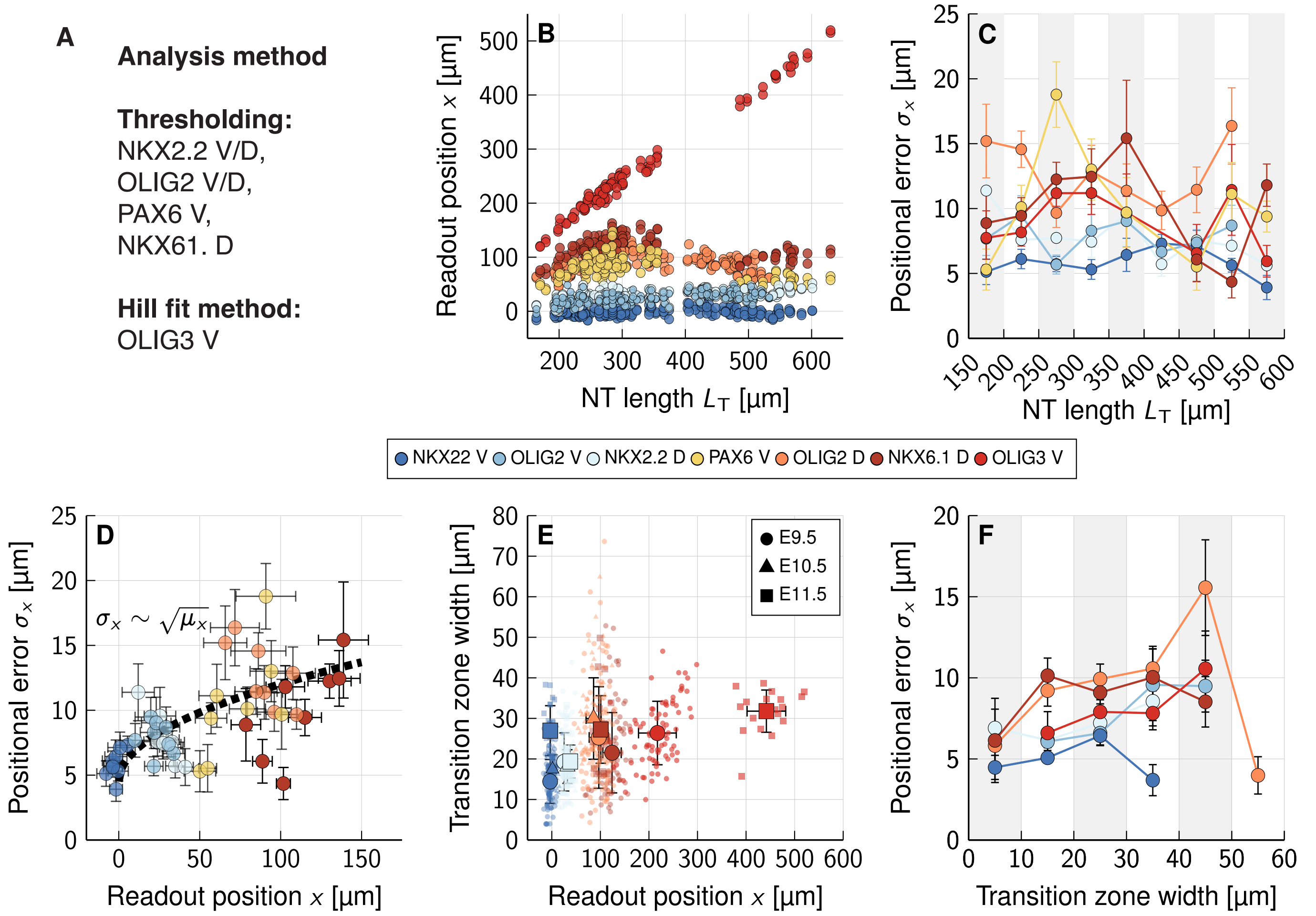


**Figure S1: Quantification of the positional error and the transition zone width in the mouse neural tube using thresholding.** **A** Analysis method for each of the domain boundaries. **B, C** Progenitor domain boundary positions (B) and positional error of the domain boundaries (C) over developmental time, as measured by increasing total NT length $L_T$. **D** Positional error vs. readout position from the ventral limit. Each point corresponds to a 50 µm bin as in (C). The black dotted line indicates the fitted theoretical scaling $\sigma_x \sim \sqrt{x}$ [9]. For fit parameters see Table S7. **E** The transition zone width (TZW) of the domain boundaries shows no systematic dependency on the readout position. **F** Transition zone widths vs. positional errors. TZWs were grouped into bins of length 10 µm. Sample numbers are listed in Tables S2, S5 and S6.

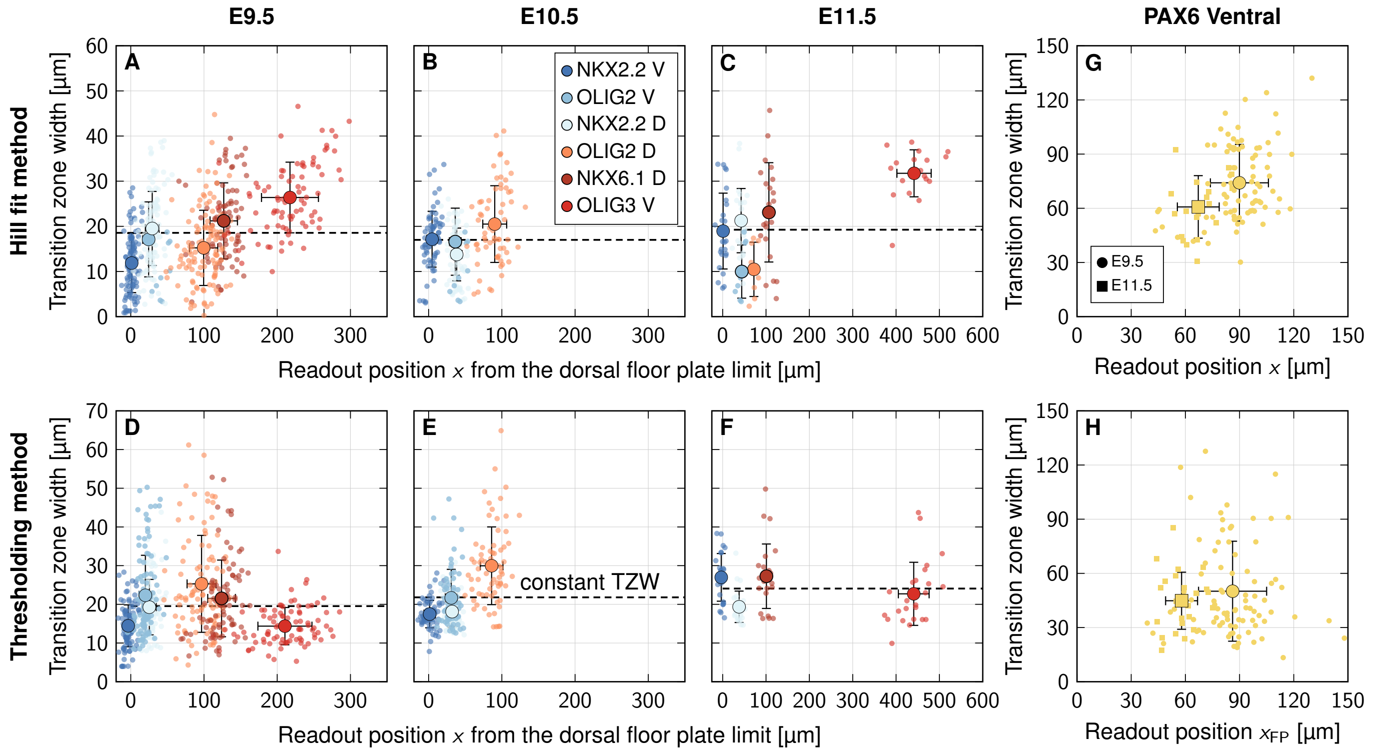


**Figure S2: Transition zone widths in the developing mouse NT. A–H** TZWs in moose embryos at developmental stages E9.5, E10.5 and E11.5. For stages E9.5 and E11.5, the ventral and dorsal NKX2.2, ventral and dorsal OLIG2, ventral PAX6, dorsal NKX6.1 and ventral OLIG3 boundaries were analysed. At stage E10.5, the dorsal NKX2.2 and dorsal OLIG2 boundaries were analysed. Small dots indicate individual NT sections, while large dots represent arithmetic means. Vertical error bars indicate standard deviations in the TZWs, horizontal error bars show standard deviations of the average readout position per developmental stage (A–F). (A–C,G) TZWs obtained via Hill fit. (D–F,H) TZWs obtained via multi-Otsu thresholding. The wide ventral PAX6 boundaries are reported separately (G,H) due to their different regulatory background. Sample numbers are listed in Tables S1 and S2.

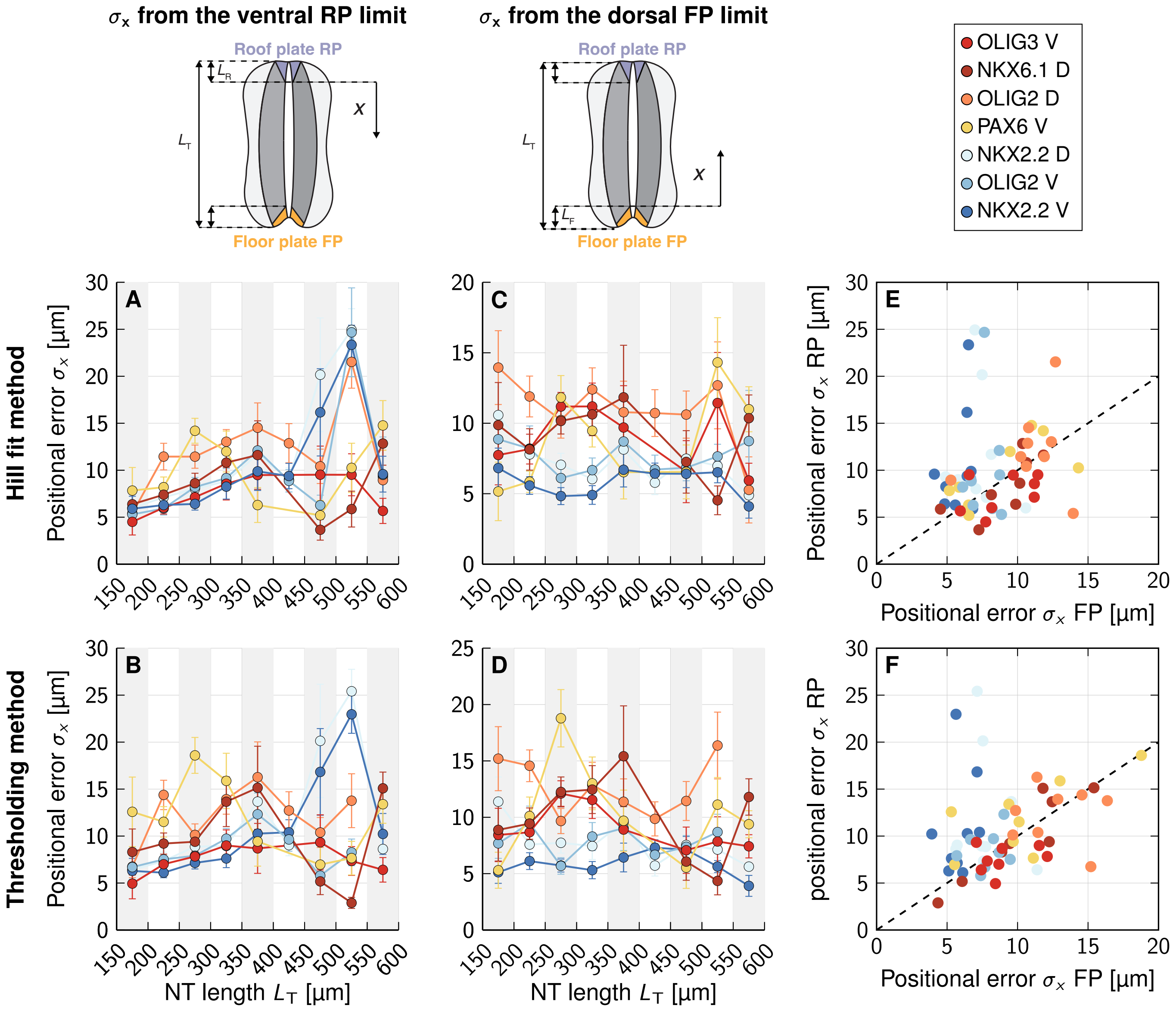


**Figure S3: Positional error of progenitor domain boundaries in the developing mouse NT. A–D** Positional errors measured with respect to the ventral limit of the roof plate (A,B) or the dorsal limit of the floor plate limit (C,D) using the Hill fit method (A,C) or the thresholding method (B,D). **E,F** Comparison of positional errors measured with respect to the FP and RP. Sample numbers are listed in Tables S3 and S5.

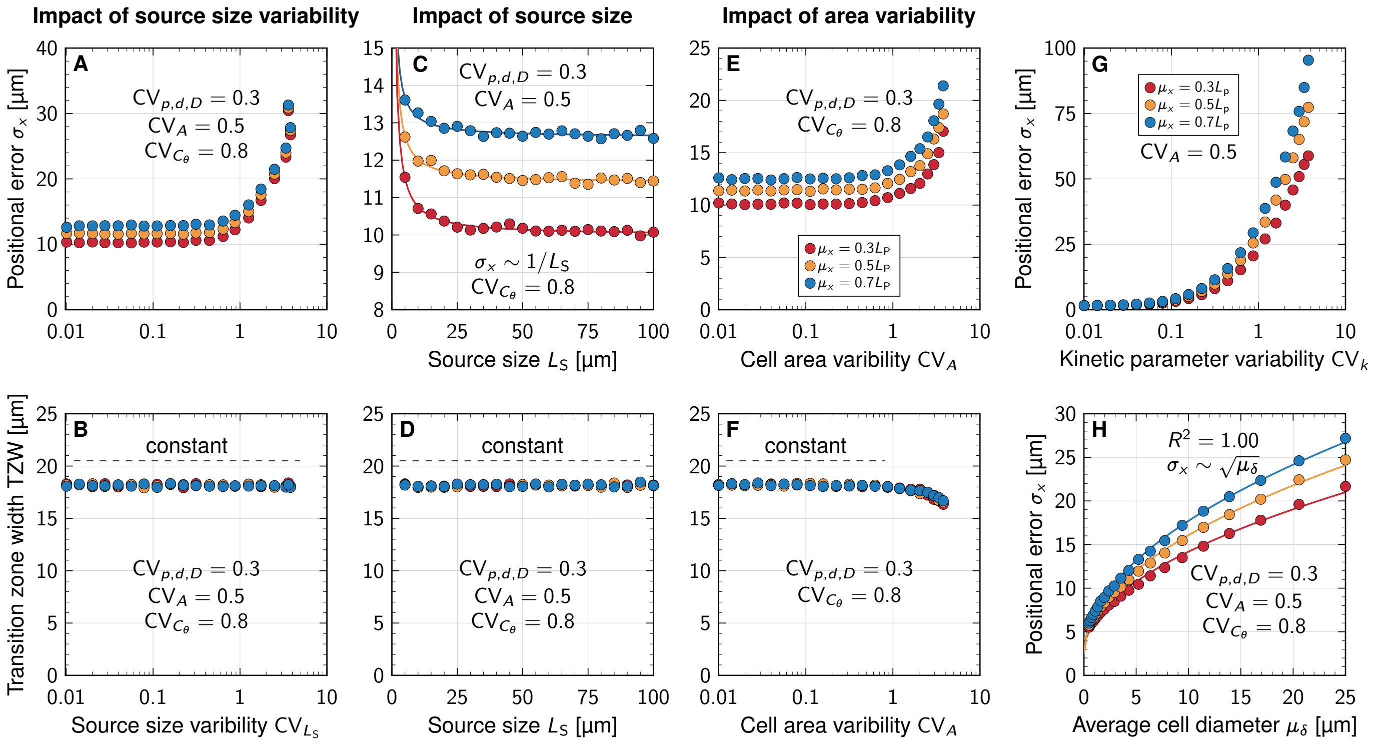


**Figure S4: Impact of cell size and different sources of noise on the TZW and positional error.** **A–D** Impact of source size variability $CV_{L_S}$ and source size $L_S$ on the positional error and TZW. **E,F** Impact of area variability $CV_A$ on the positional error and the TZW. **G** Impact of kinetic variability $CV_k$ on the positional error $\sigma_x$. **H** Scaling of the positional error with cell size. Data points are means of $N = 10^4$ independent simulations, SEM are smaller than the markers. For fit parameters see Table S7.

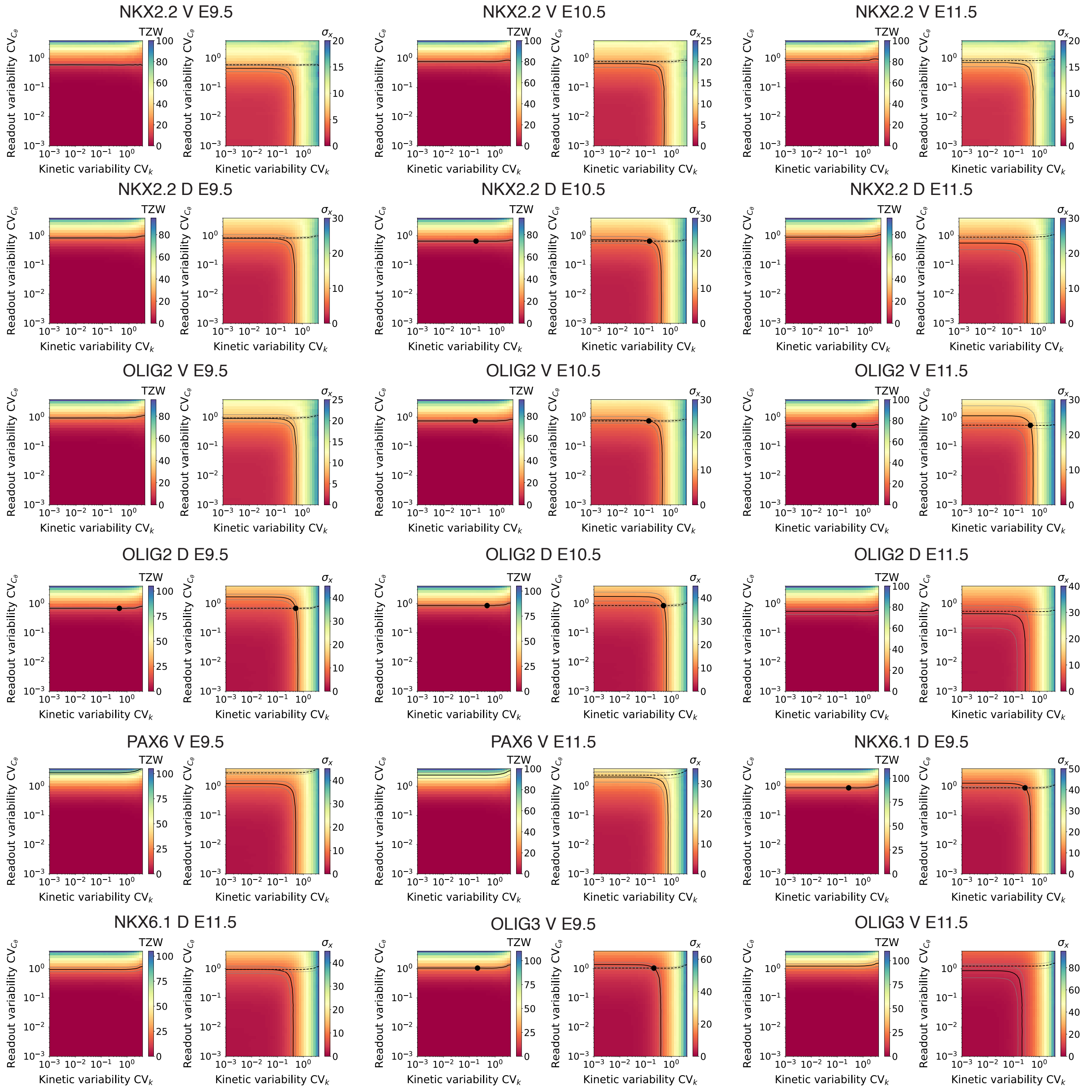


**Figure S5: Parameter screens to infer the kinetic and readout noise levels using the Hill fit approach.** Each coloured square represents the mean of $N = 10^4$ independent simulations. Contours correspond to mean TZW ± SEM or mean $\sigma_x$ ± SEM.

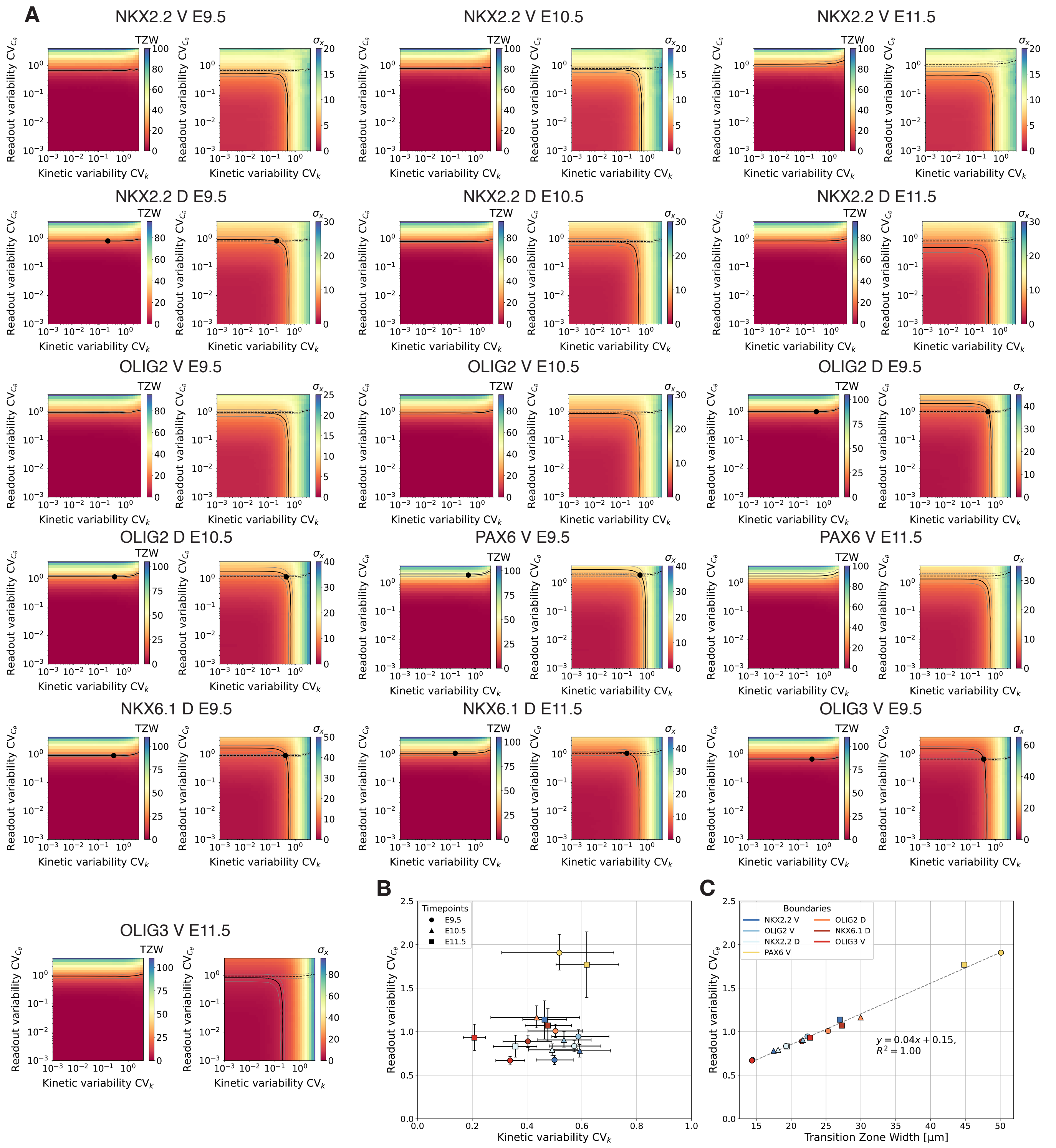


**Figure S6: Parameter screens to infer the kinetic and readout noise levels using the thresholding approach. A** Parameter screen too infer noise levels. **B** Kinetic and readout noise levels for the different domain boundaries. Each coloured square in (A) represents the mean of $N = 10^4$ independent simulations. Contours correspond to mean TZW ± SEM or mean $\sigma_x$ ± SEM. **C** Linear relationship between the measured TZW and inferred readout noise level $CV_{C_\theta}$. Error bars are omitted for clarity.

#### 2 Supplementary Tables

**Table S1:** Samples numbers for the plots in Fig. 3B,F and Fig. S2A–C,G.

| | NKX2.2 | | OLIG2 | | PAX6 | NKX6.1 | OLIG3 |
|---|---|---|---|---|---|---|---|
| Stage | V | D | V | D | V | D | V |
| E9.5 | 109 | 112 | 108 | 112 | 83 | 83 | 83 |
| E10.5 | 72 | 72 | 70 | 66 | 0 | 0 | 0 |
| E11.5 | 26 | 26 | 9 | 8 | 22 | 22 | 20 |

**Table S2:** Samples numbers for the plots in Fig. S1B,F and Fig. S2D–F,H.

| | NKX2.2 | | OLIG2 | | PAX6 | NKX6.1 | OLIG3 |
|---|---|---|---|---|---|---|---|
| Stage | V | D | V | D | V | D | V |
| E9.5 | 103 | 104 | 107 | 107 | 88 | 87 | 88 |
| E10.5 | 72 | 72 | 69 | 70 | 0 | 0 | 0 |
| E11.5 | 26 | 26 | 0 | 0 | 24 | 24 | 24 |

**Table S3:** Sample numbers for the plots in Fig. 3C,D,E and Fig. S3A,C,E.

| | NKX2.2 | | OLIG2 | | PAX6 | NKX6.1 | OLIG3 |
|---|---|---|---|---|---|---|---|
| $L_T$ [µm] | V | D | V | D | V | D | V |
| 150–200 | 7 | 8 | 8 | 8 | 4 | 4 | 4 |
| 201–250 | 40 | 40 | 37 | 40 | 20 | 19 | 20 |
| 251–300 | 34 | 36 | 35 | 36 | 41 | 42 | 41 |
| 301–350 | 24 | 24 | 24 | 24 | 12 | 12 | 12 |
| 351–400 | 8 | 8 | 8 | 8 | 4 | 4 | 4 |
| 401–450 | 26 | 26 | 26 | 26 | 0 | 0 | 0 |
| 451–500 | 24 | 24 | 22 | 19 | 4 | 4 | 4 |
| 501–550 | 34 | 34 | 20 | 19 | 6 | 6 | 6 |
| 551–600 | 6 | 6 | 4 | 3 | 10 | 10 | 8 |

**Table S4:** Sample numbers for the plot in Fig. 3G.

| | NKX2.2 | | OLIG2 | | NKX6.1 | OLIG3 |
|---|---|---|---|---|---|---|
| TZW [µm] | V | D | V | D | D | V |
| 0–10 | 53 | 35 | 41 | 39 | 9 | 0 |
| 11–20 | 111 | 95 | 87 | 87 | 35 | 22 |
| 21–30 | 36 | 64 | 44 | 41 | 39 | 38 |
| 31–40 | 7 | 16 | 11 | 13 | 16 | 34 |
| 41–50 | 0 | 0 | 0 | 3 | 0 | 5 |

**Table S5:** Sample numbers for the plots in Fig. S1C,D and Fig. S3B,D,F.

| | NKX2.2 | | OLIG2 | | PAX6 | NKX6.1 | OLIG3 |
|---|---|---|---|---|---|---|---|
| $L_T$ [µm] | V | D | V | D | V | D | V |
| 150–200 | 8 | 8 | 8 | 8 | 4 | 4 | 4 |
| 201–250 | 37 | 38 | 37 | 37 | 22 | 22 | 22 |
| 251–300 | 36 | 36 | 34 | 34 | 44 | 43 | 44 |
| 301–350 | 18 | 18 | 24 | 24 | 12 | 12 | 12 |
| 351–400 | 8 | 8 | 8 | 8 | 4 | 4 | 4 |
| 401–450 | 26 | 26 | 25 | 25 | 0 | 0 | 0 |
| 451–500 | 24 | 24 | 22 | 22 | 4 | 4 | 4 |
| 501–550 | 34 | 34 | 14 | 15 | 6 | 6 | 6 |
| 551–600 | 6 | 6 | 0 | 0 | 12 | 12 | 12 |

**Table S6:** Sample numbers for the plot in Fig. S1F.

| | NKX2.2 | | OLIG2 | | NKX6.1 | OLIG3 |
|---|---|---|---|---|---|---|
| TZW [µm] | V | D | V | D | D | V |
| 0–10 | 21 | 10 | 2 | 5 | 5 | 14 |
| 11–20 | 123 | 111 | 87 | 42 | 41 | 76 |
| 21–30 | 46 | 64 | 52 | 68 | 42 | 14 |
| 31–40 | 7 | 12 | 18 | 32 | 10 | 2 |
| 41–50 | 0 | 0 | 12 | 19 | 7 | 0 |
| 51–60 | 0 | 0 | 0 | 4 | 0 | 0 |

**Table S7:** Summary of the fit functions and their parameters. All parameters are in micrometers.

| **Figure** | **Model** | **Legend entry** | $\boldsymbol{a}$ | **SE**($\boldsymbol{a}$) | $\boldsymbol{b}$ | **SE**($\boldsymbol{b}$) |
|---|---|---|---|---|---|---|
| 3E | $a\sqrt{\mu_x} + b$ | — | 0.45 | 0.07 | 5.14 | 0.53 |
| 4E | $a\ln(b/\mu_\delta)$ | $\mu_x = 0.3L_\mathrm{P}$ | 11.6 | 0.1 | 25.3 | 0.7 |
| | | $\mu_x = 0.5L_\mathrm{P}$ | 11.6 | 0.1 | 26.2 | 0.7 |
| | | $\mu_x = 0.7L_\mathrm{P}$ | 7.5 | 1.1 | 25.0 | 0.7 |
| S1D | $a\sqrt{\mu_x} + b$ | — | 0.60 | 0.16 | 5.19 | 1.28 |
| S4C | $a/L_\mathrm{S} + b$ | $\mu_x = 0.3L_\mathrm{P}$ | 7.4 | 1.6 | 10.0 | 0.1 |
| | | $\mu_x = 0.5L_\mathrm{P}$ | 8.1 | 1.5 | 11.3 | 0.1 |
| | | $\mu_x = 0.7L_\mathrm{P}$ | 7.6 | 1.6 | 12.5 | 0.1 |
| S4H | $a\sqrt{\mu_\delta} + b$ | $\mu_x = 0.3L_\mathrm{P}$ | 3.66 | 0.07 | 2.53 | 0.17 |
| | | $\mu_x = 0.5L_\mathrm{P}$ | 4.33 | 0.05 | 2.31 | 0.12 |
| | | $\mu_x = 0.7L_\mathrm{P}$ | 4.79 | 0.05 | 2.35 | 0.13 |

**Table S8:** Hill coefficient by stage and domain (mean ± SD; SE in parentheses).

| Domain | Side | E9.5 | E10.5 | E11.5 |
|---|---|---|---|---|
| PAX6 | V | 0.51 ± 0.18 (0.02) | — | 0.99 ± 0.31 (0.07) |
| NKX2.2 | V | 4.99 ± 6.07 (0.58) | 3.94 ± 3.32 (0.39) | 3.66 ± 1.98 (0.39) |
| NKX2.2 | D | 2.22 ± 1.76 (0.17) | 4.78 ± 2.66 (0.31) | 2.94 ± 1.23 (0.24) |
| OLIG2 | V | 2.58 ± 1.81 (0.17) | 4.02 ± 2.70 (0.32) | 7.92 ± 5.05 (1.68) |
| OLIG2 | D | 4.15 ± 9.85 (0.93) | 3.21 ± 2.04 (0.25) | 8.45 ± 7.29 (2.58) |
| NKX6.1 | D | 2.01 ± 1.21 (0.13) | — | 3.50 ± 3.01 (0.64) |
| OLIG3 | V | 1.44 ± 0.55 (0.06) | — | 1.81 ± 0.45 (0.10) |

#### Supplementary Notes

##### Derivation of the previously proposed transition zone width at low molecule numbers

The width of the transition zone has previously been related to the noisiness of morphogen gradients assuming low receptor occupancy [8]. We recapitulate this idea here to derive Eq. 3, which we showed does not apply in the developing neural tube.

The argument is that at sufficiently low molecule numbers, the variability of the signal the cells are exposed to would be dominated by "binding noise", as ligand-receptor binding becomes a Poisson process. In this stochastic setting, the concentration threshold $C_\theta$ is reached at multiple different readout positions $x_\theta$ inside the patterning domain (Fig. S7), leading to a salt-and-pepper expression pattern. One may then quantify the transition zone width as twice the standard deviation of the salt-and-pepper readout positions, $\mathrm{TZW} = 2\sigma_x$, with $\sigma_x = \mathrm{SD}[x_\theta]$, at least in proportion. One may then use linear error propagation to express the positional variation in terms of molecular fluctuations:

$$\mathrm{TZW} = 2\sigma_x \approx 2\left|\frac{\partial C}{\partial x}\right|^{-1}\sigma_C. \tag{9}$$

Here, $\sigma_C(x) = \mathrm{SD}[C(x)]$ quantifies that signalling variability locally at a given point in the tissue, $x$. Assuming that the morphogen concentration is exponential (Eq. 1) aside from its local variability, it follows that

$$\frac{\partial C}{\partial x}(x) = -\frac{C(x)}{\lambda},$$

such that $|\partial C/\partial x|^{-1} = \lambda/C(x)$. The transition zone width then reads [8]

$$\mathrm{TZW} \approx 2\lambda \mathrm{CV}_C, \tag{10}$$

where $\mathrm{CV}_C = \sigma_C/\mu_C$ is the coefficient of variation of the local morphogen concentration. Assuming a Poissonian nature of the signal exposure implies that $\mathrm{CV}_C$ would be inversely proportional to the square root of the local morphogen number, since $\mathrm{CV} = 1/\sqrt{\mu}$ for the Poisson distribution. The width of the transition zone should then increase exponentially with its distance from the source:

$$\mathrm{TZW} \propto \lambda \exp\left[\frac{x}{2\lambda}\right], \tag{11}$$

which we do not observe in the mouse NT (Fig. 3F).

##### Derivation of the TZW using Hill functions

To find an analytical solution for the TZW, we plug the exponential morphogen gradient $C(x) = C_0 \exp[-x/\lambda]$ into the Hill equation,

$$H(x) = \frac{1}{1 + (K/C(x))^n}, \tag{12}$$

which we assume to underlie the regulatory readout by the cells. We then invert the Hill equation and determine the positions $x_i$ and $x_j$ where it reaches two distinct thresholds $H_i = H(x_i) \in (0,1)$, $i = 1, 2$. In line with our definition of the transition zone, one may use, for instance, $H_1 = 0.9$ and $H_2 = 0.1$. Solving for the distance between them yields

$$\begin{aligned}\mathrm{TZW} = x_2 - x_1 &= \frac{\lambda}{n}\ln\left[\frac{1/H_2 - 1}{1/H_1 - 1}\right] \\ &= \frac{\lambda}{n}\big(\mathrm{logit}[H_1] - \mathrm{logit}[H_2]\big).\end{aligned} \tag{13}$$

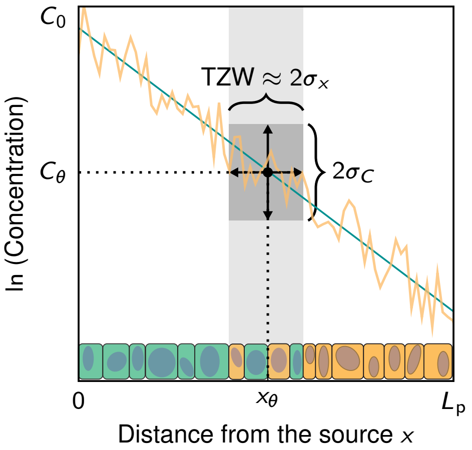


**Figure S7: Approximation of the transition zone width from noisy morphogen gradients.** The transition zone width can be estimated using the average morphogen gradient profile $\mu_C$ and its variability $\sigma_C$ through linear error propagation.

Note that Eq. 13 is equivalent to the formula [7] for the width $\Delta x$ of a domain defined by two concentrations thresholds $C_1$ and $C_2$,

$$\Delta x = x_2 - x_1 = \lambda \ln\left[\frac{C_1}{C_2}\right] \tag{14}$$

if we set $C_i \propto \sqrt[n]{1/H_i - 1}$, $i = 1, 2$.

##### The sample standard deviation of the log-normal distribution is a biased estimator

The log-normal distribution is characterised by two parameters $\mu$ and $\sigma$. To produce a distribution with desired mean $\mu_X$ and $\sigma_X$ one uses the relationships

$$\mu = \ln\left[\frac{\mu_X}{\sqrt{1 + \sigma_X^2/\mu_X^2}}\right] = \ln\left[\frac{\mu_X}{\sqrt{1 + \mathrm{CV}_X^2}}\right], \tag{15}$$

and

$$\sigma = \sqrt{\ln\left[1 + \frac{\sigma_X^2}{\mu_X^2}\right]} = \sqrt{\ln\left[1 + \mathrm{CV}_X^2\right]}. \tag{16}$$

For large coefficients of variation, the log-normal distribution becomes broad and heavy-tailed (Fig. S8A). This characteristic leads to an underestimation of the true CV by the sample CV with small populations (Fig. S8B). In this study, we report the sample CV, calculated from the sample standard deviation $s$ and the sample mean $\bar{x}$,

$$\widehat{\mathrm{CV}} = \frac{s}{\bar{x}}. \tag{17}$$

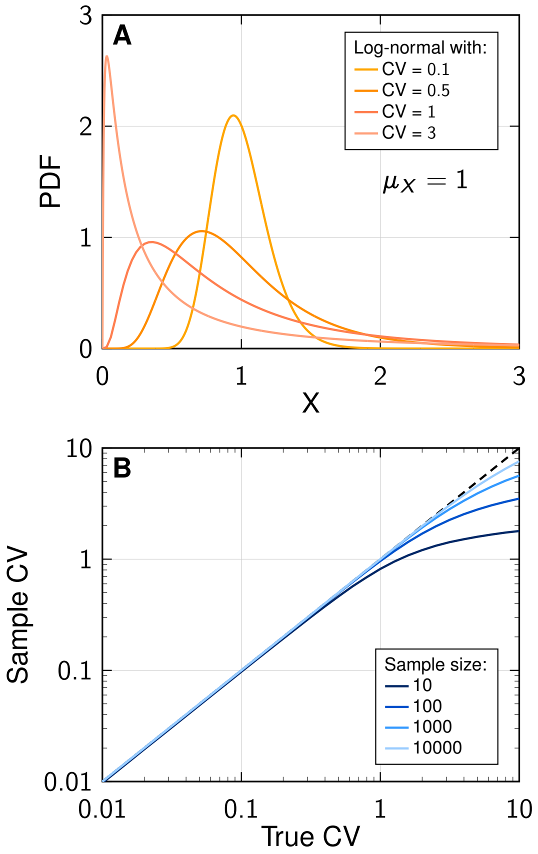


**Figure S8: Estimation of the coefficient of variation from log-normally distributed data.** **A** Probability density function (PDF) of different log-normal distributions with unit mean. **B** Deviation of true and sample CV values for finite populations.
